# Noise ceiling on the crossvalidated performance of reweighted models of representational dissimilarity: Addendum to Khaligh-Razavi & Kriegeskorte (2014)

**DOI:** 10.1101/2020.03.23.003046

**Authors:** Katherine R. Storrs, Seyed-Mahdi Khaligh-Razavi, Nikolaus Kriegeskorte

## Abstract

An error was made in including noise ceilings for human data in Khaligh-Razavi and Kriegeskorte (2014). For comparability with the macaque data, human data were averaged across participants before analysis. Therefore the noise ceilings indicating variability across human participants do not accurately depict the upper bounds of possible model performance and should not have been shown. Creating noise ceilings appropriate for the fitted models is not trivial. Below we present a method for doing this, and the results obtained with this new method. The corrected results differ from the original results in that the best-performing model (weighted combination of AlexNet layers and category readouts) does not reach the lower bound of the noise ceiling. However, the best-performing model is not significantly below the lower bound of the noise ceiling. The claim that the model “fully explains” the human IT data appears overstated. All other claims of the paper are unaffected.

---

Noise ceilings should not have been included in the plots of Khaligh-Razavi and Kriegeskorte (2014) showing correlations between models and human inferior temporal (hIT) data (Figures 2A and 7, and Supplementary Figures 2, 3 and 12), as they were not statistically appropriate. The noise ceilings were calculated as implemented in the RSA Toolbox (Nili et al., 2014), by computing the average correlation of each participant’s representational dissimilarity matrix (RDM) with the group average RDM, either including (upper bound) or excluding (lower bound) that participant’s data. This provides the upper bound on the possible model performance when model performance is calculated by correlating the model RDM separately with each participant’s RDM and averaging these correlations. However, in the paper, all model evaluations were performed against the single group-average RDM, for better comparability with the macaque RDM (which averages over N=2 macaques). Inter-subject variance therefore does not limit possible model performance, and the correct upper bound is 1. The figures for the macaque data do not contain noise ceilings for this reason, and neither should those for the human data.

The removal of noise ceilings from Figures 2 and 7 leaves unsupported the claim that reweighted features from a deep supervised model fully explain hIT data. Here we present a new analysis to assess whether this claim holds. One approach would be to estimate the lower bound of the noise ceiling correctly for the bar graphs shown in the paper. However, this is not trivial and requires distributional assumptions, which we would like to avoid. We therefore instead take the approach of Nili et al. (2014) for the noise ceiling and plot the group-average RDM correlation for each model (rather than the correlation with the group-average RDM). The Nili et al. (2014) approach to estimating model performance (group-average of single-subject RDM correlations) requires a separate RDM prediction for each subject. A complication is that some of the models have parameters to be fitted, requiring crossvalidation across images and subjects for unbiased estimates of model performance.

Below we describe a novel and flexible analysis method for fitted RDM models. On each fold of crossvalidation, the training and test set is based on a nonoverlapping set of stimuli and a nonoverlapping set of subjects. Each model is fitted to the training set and evaluated on the test set (different subjects and stimuli). The lower and upper bounds of the noise ceiling are computed in the same crossvalidation loop. The performance estimates and noise-ceiling estimates (RDM correlations) are averaged across folds. The entire crossvalidation procedure can be repeated for many bootstrap samples of stimuli and subject. Here the bootstrapping was limited to stimuli, because there were only four subjects in the dataset.

## Method for estimating model performances and noise-ceiling bounds

The performance of each model, and the lower and upper bounds of the noise ceiling, were computed concurrently, using the following procedure:

1. For 1000 stimulus-bootstrap samples:

a. For 20 stimulus-crossvalidation folds:

i. Randomly assign 8 unique stimuli present in this bootstrap sample to be test stimuli. As a result, the test set always consists of data from exactly 8 unique images, and typically contains repetitions of some of them. Data from the same image never appears in both training and test sets, whether or not it is repeatedly present in the bootstrap sample.
ii. For 4 subject-crossvalidation folds (leaving out each subject in turn):

1. Average the data RDMs across training subjects for the training images. Using the resulting training RDM, fit the weights of the parameterized models with non-negative least squares regression (e.g., in Figure 1 below, the components are the 7 Alexnet layer RDMs and 3 SVM-remixed RDMs).
2. For each model, create a predicted RDM for the test images (using fitted weights for parameterized models). Calculate each model’s performance as the correlation between the model-predicted RDM and the test RDM (the RDM for the test images in the test subject).
3. At the same time, calculate the upper and lower bounds of the noise ceiling by taking the correlation between the test RDM and the average test-image RDM for either all 4 subjects (upper bound) or the 3 training subjects (lower bound).
iii. For each test subject, we now have an estimate of each model’s performance and of the upper and the lower bounds of the noise ceiling. Average these to create a single estimate of each model’s performance and the noise ceiling bounds, for this stimulus-crossvalidation sample.
b. At the end of the 20 stimulus-crossvalidation folds, we have 20 estimates for each model’s performance and 20 estimates of the upper and the lower bounds of the noise ceiling. Average these to create a single estimate of each of these measures, for this bootstrap sample.
2. At the end of this procedure, we have a distribution of 1000 estimates of each model’s performance and of the upper and lower bounds of the noise ceiling, bootstrapped over the population of image stimuli used in the experiment.

**Figure 1:**
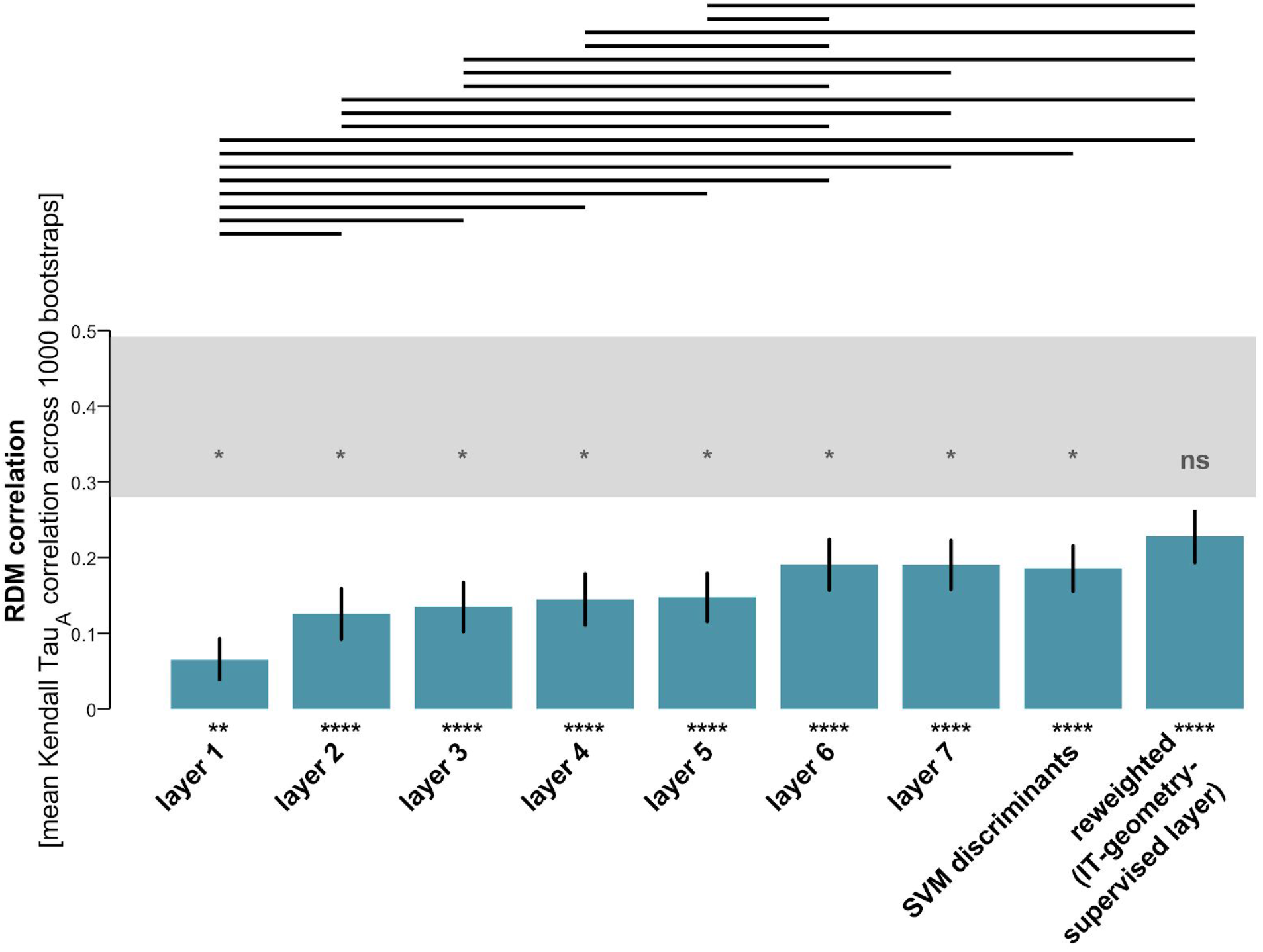
The strongly supervised deep network, with features remixed and reweighted, best explains the IT data. The bars show the Kendall-*τ_A_* RDM correlations between the layers of the strongly supervised deep convolutional network and human IT. The error bars are standard errors of the mean estimated by bootstrap resampling of the stimuli. Asterisks indicate significant RDM correlations (p < 0.05: *, p < 0.01: **, p < 0.001: ***, p < 0.0001: ****; inference by bootstrap resampling of the stimuli). As we ascend the layers of the deep network, model RDMs explain increasing proportions of the variance of the hIT RDM. The noise ceiling (gray bar) indicates the expected performance of the true model (given the noise and intersubject variability). The true model here refers to the unknown process that actually generated the data. The upper and lower edges of the noise ceiling bar are upper and lower bound estimates on the performance of the true model. None of the layers of the deep network reaches the noise ceiling. Remixing the features of layer 7 using linear SVMs to strengthen the categorical divisions creates representations composed of three discriminants: animate/inanimate, face/nonface, and body/nonbody. The “SVM discriminants” bar indicates the performance of a uniformly-weighted combination of the three SVM discriminants. Reweighting the model layers and the three discriminants using the cross-validation procedure outlined in the text yields a representation that explains the hIT geometry best. A horizontal line over two bars indicates that the two models perform significantly differently (inference by bootstrap resampling of the stimulus set). Multiple testing across the many pairwise comparisons is accounted for by controlling the FDR at 0.05. A grey asterisk above the lower bound of the noise ceiling indicates that a model is significantly below the lower bound, controlling the FDR at 0.05 (alternatively, ns for not significant). All models are significantly below the lower bound of the noise ceiling, except for the hIT-geometry-supervised deep model. This figure supplements Figure 7 in the original paper.

**Figure 2:**
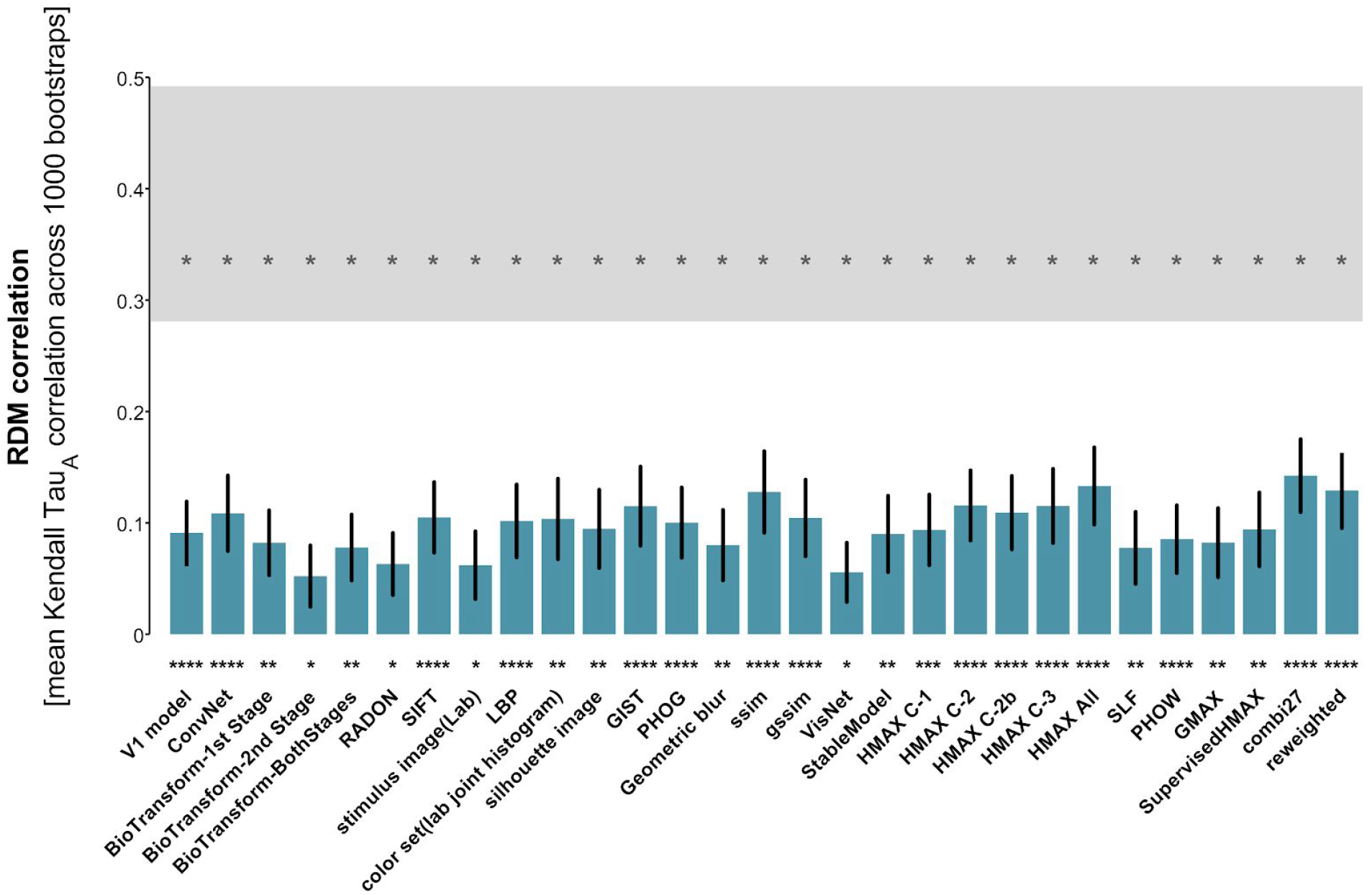
The not-strongly-supervised models fail to fully explain the IT data. The bars show the Kendall-*τ_A_* RDM correlations between the not-strongly-supervised models and human IT. The error bars are standard errors of the mean estimated by bootstrap resampling of the stimuli. Asterisks indicate significant RDM correlations (p < 0.05: *, p < 0.01: **, p < 0.001: ***, p < 0.0001: ****). Conventions are the same as for Figure 1. The “combi27” model is a uniformly-weighted combination of the other 27 models, and the final “reweighted” model is a combination of all 28 previous models using weights estimated by fitting to the human IT data. Performance of each model was estimated using the cross-validation procedure outlined in the text. As reported in the original paper, the fitted combination model performs slightly worse than the uniformly weighted model, due to overfitting. None of the not-strongly-supervised models reaches the noise ceiling. This figure supplements Figure 2A in the original paper.

Figure 1 shows the performance of the deep supervised neural network model from Khaligh-Razavi and Kriegeskorte (2014) in explaining their hIT data, calculated using the above procedure (cf. Figure 7 in the original paper), and Figure 2 does the same for the 27 shallow and not-strongly-supervised models (cf. Figure 2A in the original paper).

## Conclusions

The noise ceilings shown in figures presenting human fMRI results in Khaligh-Razavi and Kriegeskorte (2014) are inappropriate, and should not have been included. A novel analysis method enables us to estimate noise ceilings for fitted and unfitted models, and to inferentially compare the models to each other and to the lower bound of the noise ceiling. The best-performing model no longer reaches the noise ceiling, but is statistically indistinguishable from its lower bound. The main claims of the paper are therefore unaffected.

